# Microbial Diversity and Function Linked to Carbon Cycling in Mangrove Sediments

**DOI:** 10.64898/2026.05.13.724760

**Authors:** Nabilah Khairi, Muhammad Zarul Hanifah Md Zoqratt, Nur Hidayah, Bahaa Abdella, Muhammad Azamuddeen Mohammad Nasir, Geok Yuan Annie Tan, Phaik-Eem Lim, Wan Syaidatul Aqma, Ahmad Aldrie Amir, Mohammad Rozaimi, Nur Hazlin Hazrin-Chong

## Abstract

Microbial communities are central to the biogeochemical cycling of nutrients, critically shaping ecosystem functioning and influencing climate change mitigation. Mangrove ecosystems are among the most important global carbon sinks that enable large amounts of carbon to be sequestered and stored. However, gaps persist in understanding the fundamental aspects of microbial-driven carbon cycling in these environments. This research explores the microbial taxonomic and functional diversity related to carbon cycling in selected tropical mangrove sediments across various locations and depths. Sequencing data analyses based on the 16S rRNA gene revealed distinct microbial community composition but conserved predicted functions across the different mangrove locations. Depth was a strong influence on the functional composition, with carbon-related pathways and metabolic strategies differing between top and bottom sediments. Putative functional gene abundance analyses revealed that carbon fixation processes were among the top carbon-related pathways, suggesting the key role of mangrove microbial communities in sustaining long-term carbon storage. Within these communities, Desulfobacterota appeared as a primary contributor to carbon fixation, while Chloroflexota played a significant role in carbon metabolism and methane cycling. Co-occurrence network analyses also revealed that these microbial groups were among the keystone taxa in mangrove sediments. Our study adds on to the body of knowledge on the mangrove microbiome and their carbon metabolic processes, which helps to improve strategies for managing and leveraging these vital carbon sinks.

## 1. Introduction

Mangrove ecosystems represent among the most productive and ecologically significant biomes on Earth (Donato *et al*., 2011). Despite covering only approximately 0.1% of the global continental surface, mangroves store up to 15% of the organic carbon (OC) contained in sediments (Duarte *et al*., 2013). Mangroves possess the largest carbon pool of 6.5 billion tons out of 11.5 billion tons in blue carbon ecosystems, which include seagrass meadows and salt marshes (Siikamäki *et al*., 2012). This highlights the vital role of mangroves in climate regulation through carbon sequestration and long-term storage, above- and belowground (Atwood *et al*., 2017; Macreadie *et al*., 2021). Mangrove sediments constitute approximately 85% of the total ecosystem’s carbon storage, reaching 4.4 to 11.7 pentagram of carbon (Pg C), and contribute around 26.1 ± 6.3 teragram of Carbon (Tg) C for long-term burial (Kauffman *et al*., 2020). The role of mangrove sediments as effective carbon sinks can be attributed to their high sedimentation rates, waterlogged conditions, and limited oxygen, which slow decomposition processes (Alongi, *et al*., 2012; Atwood *et al*., 2017; Choudhary *et al*., 2024).

Microbial communities play a central role in regulating the fate of OC through diverse metabolic pathways. In particular, the role of these microorganisms facilitates carbon cycling by decomposing organic matter, transforming nutrients and stabilising carbon through biogeochemical processes (Kristensen *et al*., 2008; Trevathan-Tackett *et al*., 2017). In addition to microbial by-products and necromass that contribute to stabilised carbon fractions, microbial communities contribute to carbon storage by selectively degrading labile compounds, leading to the accumulation of more recalcitrant organic matter that persists and is eventually buried in deeper sediment layers. Meanwhile, through microbial respiration, organic matter is remineralised, which releases carbon as CO_2_, bicarbonate and carbonate into porewater (Garritano *et al*., 2022). Alternative respiration pathways, e.g., sulfate reduction and methanogenesis in deeper sediments, also contribute to carbon turnover and influence whether carbon is stored or emitted as greenhouse gases.

Although previous studies have established the importance of microbial processes in mangrove carbon cycling, our understanding remains incomplete. In particular, the specific microbial taxa and metabolic pathway that support carbon sequestration in mangrove sediments are still poorly defined. This knowledge gap is especially pronounced in Southeast Asia mangroves, which are among the most carbon-dense globally yet remain underrepresented in microbial ecology research. Most existing studies focus on carbon stocks, burial rates and organic carbon dynamics in mangrove ecosystems (Alongi, 2014; Bouillon *et al*., 2008), while microbial contributions especially carbon fixation, metabolic redundancy, and depth-dependent functional shifts have received far less attention. Furthermore, reviews of mangrove microbiology have emphasised that the phylogenetic and functional diversity of mangrove microorganisms insufficiently explored despite their crucial roles in nutrient cycling and ecosystem functioning (Thatoi *et al*., 2013; Lai *et al*., 2022). These knowledge gaps limit our ability to fully explain why mangrove sediments accumulate such large amounts of carbon and importantly, constrain predictions of how these carbon sinks will respond to rapid environmental changes.

To address these knowledge gaps, we explored the taxonomic and functional profiles of microbial communities related to carbon cycling across spatial and vertical gradients of mangrove sediments in Malaysia. We found that the mangrove sediment microbiome exhibits site-specific compositions, influenced by physicochemical factors like pH, salinity, and sediment type. Despite this taxonomic variability, metabolic functions, including related to carbon, remain conserved across locations, suggesting functional redundancy in mangrove sediments. Depth was shown as a strong influence driving these microbial functions. Key microbial groups associated with carbon fixation, including Desulfobacterota and Chloroflexota, were found to be among the keystone taxa in these mangrove sediments. Together, our results highlight the microbial communities and their contributions to carbon cycling in mangrove sediments that govern this vital blue carbon sink.

## 2. Materials and Methods

### 2.1 Site description and sample collection

Mangrove sediment cores were collected in March - September 2022 from three different locations in Peninsular Malaysia: Seri Buat Island (SB), Sg Pulai Estuary (PL), and Kuala Selangor (KS) (Fig. 1), using a steel auger (120 cm length, 5.5 cm diameter). At each site, three to five sediment cores were extracted, with sampling points spaced at least 10 meters apart. Each core was sectioned into three depth intervals: 0–10 cm (top layer, T), 40–50 cm (middle layer, M), and 80–100 cm (bottom layer, B). In addition to mangrove sediments, seagrass sediment samples were collected from Merambong (MB) to provide a comparison between different ecosystem types in selected analyses. The dominant mangrove species at each study site were recorded during sampling. PL was primarily dominated by Rhizophora apiculata. SB consisted of a mixed Rhizophora-Bruguiera forest and KS was mainly dominated by Bruguiera parviflora. All sediment samples were stored at 4 °C during transportation and transferred to −80 °C for long-term storage before further analyses.

**Fig 1.**
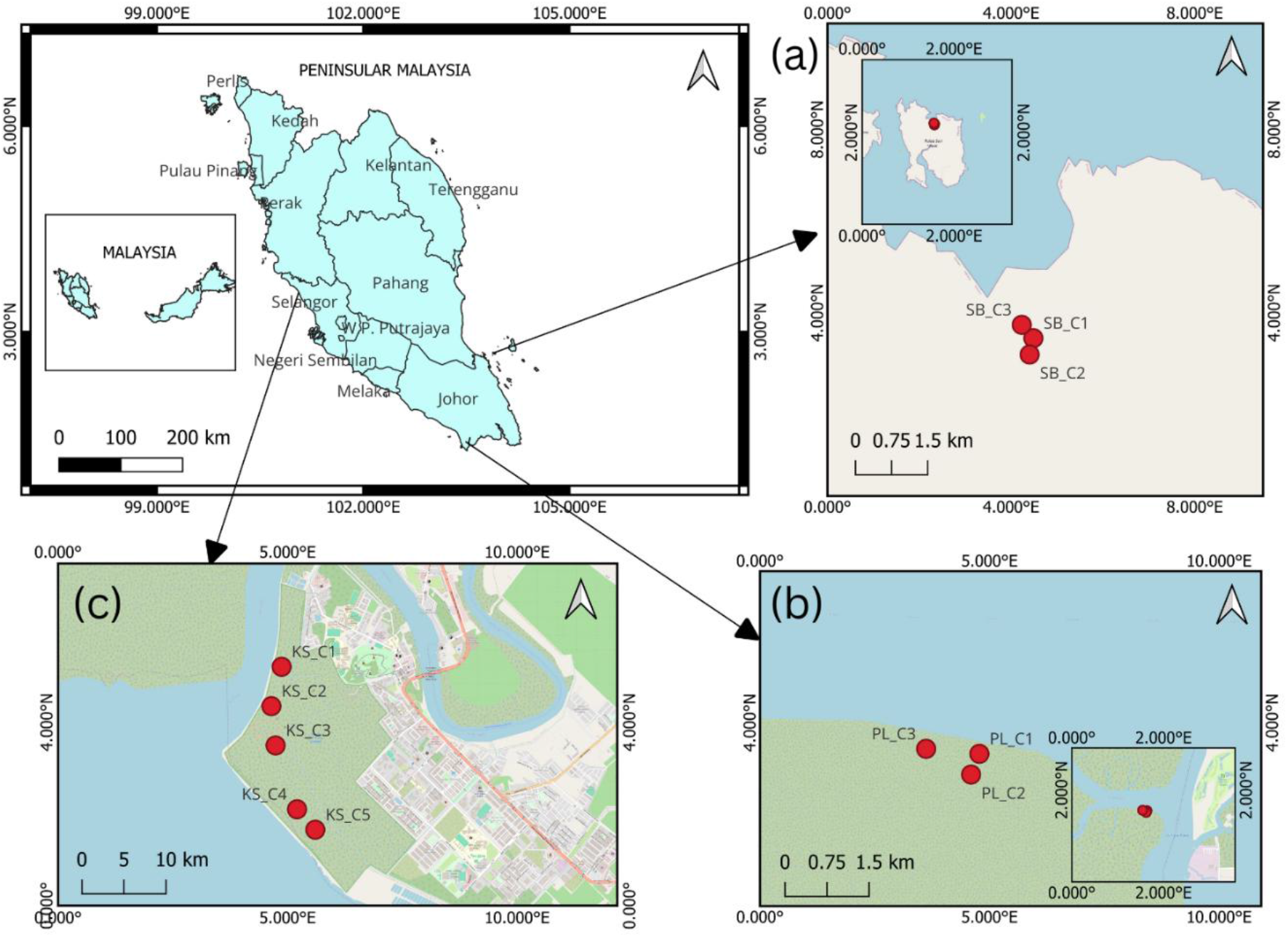
Sample sites from three different locations in Peninsular Malaysia: (a) Seri Buat Island (SB) with an inlet showing the whole island of Pulau Seri Buat, (b) Sg Pulai Estuary (PL), and (c) Kuala Selangor Nature Park (KS). This map was generated by using QGIS version 3.8 (http://www.qgis.org).

### 2.2 Physicochemical and biological analyses of mangrove sediments

Baseline physicochemical and biological analyses were conducted on sediment samples collected from the three mangroves locations. Sediment temperatures were measured in situ immediately after core extraction at multiple depths (0–10 cm, 40–50 cm, and 80–100 cm) using a thermometer probe. For laboratory measurements, soil samples were first homogenised, and a soil slurry was prepared by mixing sediment with deionized water at a 1:2.5 (w/v) ratio. The pH and salinity of the slurry were measured using a HANNA HI98194 Multiparameter Water Quality Meter (Hanna Instruments, USA). Measurements were performed in triplicate for each sample.

Total organic carbon (TOC) and total nitrogen (TN) were determined by the loss on ignition method following Rozaimi *et al*. (2017). Bulk density was calculated from oven-dried (60 °C) 50 g subsamples, which were then ground using a ball mill grinder (Retsch GmbH, Haan, Germany). Calcium carbonate content was analysed by sequential combustion in a muffle furnace following the method of Heiri *et al*. (2001). Organic carbon (OC) and stable isotope ratios (δ^13^C and δ^15^N) were analysed using a continuous flow isotope ratio mass spectrometer at the Stable Isotope Core Lab, Washington State University (USA). Particle size distribution was measured using a Laser Scattering Particle Size Distribution Analyzer (HORIBA Scientific, Japan) following the method of Callesen *et al*. (2018), after pretreatment with hydrogen peroxide to remove organic matter.

Flow cytometry quantified cell abundance following a modified protocol of Duhamel and Jacquet (2006). Sediment samples were fixed with glutaraldehyde, stained with SYBR Green, and analysed using a BDTM FACSCanto II flow cytometer (BD Biosciences, USA) with fluorescence microspheres (BDTM Liquid Counting Beads, 1 μm) as internal standards. Data analysis was performed using FACS Diva software version 6.1.3 (BD Biosciences).

### 2.3 DNA extraction, 16S rRNA gene amplification, and sequencing

Genomic DNA was extracted from 0.5 g of each sediment sample collected from the mangrove and seagrass locations using the SPINeasy DNA Kit for Soil (MP Biomedicals, USA) following the manufacturer’s protocol. Extraction blanks were included as negative controls. DNA quality and quantity were assessed using a NanoDrop 2000 Spectrophotometer (Thermo Fisher Scientific, USA) and 1% agarose gel electrophoresis, and only samples with concentrations above 20 ng/µL and a 260/280 ratio of approximately 1.8 were used for amplification.

The V3–V4 regions of the 16S rRNA gene were amplified using primers with partial Illumina Nextera adapter sequences (Klindworth *et al*., 2013) following the Illumina 16S Metagenomic Library Preparation Protocol. PCR products were verified on 1.5% (w/v) agarose gels and purified with the PrimeWay Gel Extraction (Biobasic, Canada). Amplicons were barcoded with unique dual-index primers to enable multiplexing, quantified, pooled in equimolar concentrations, and prepared for sequencing on the Illumina MiSeq platform. Samples from PL, SB, and MB were sequenced with a 2 × 300 bp paired-end configuration, while KS samples were sequenced with a 2 × 250 bp configuration. To ensure data consistency across platforms, a ZymoBIOMICS^™^ Microbial Community Standard (Zymo Research, USA) was included as an internal control in both sequencing runs.

### 2.4 Amplicon sequence data processing and statistical analyses

Amplicon sequence data were first subjected to quality trimming using Trimmomatic (Bolger *et al*., 2014) based on parameters as follows: TRAILING:25, MINLEN:250. Primers were later trimmed off using Cutadapt (Martin, 2011); untrimmed sequences were discarded. Paired-end sequences were merged using vsearch fastq_mergepairs (Rognes *et al*., 2016). The merge sequences were then denoised to generate amplicon sequence variants (ASVs) sequences using DADA2 (Callahan *et al*., 2016) within the QIIME 2 pipeline (Bolyen *et al*., 2019). The taxonomic classification of ASVs was performed using a q2-feature-classifier (Bokulich *et al*., 2018) trained on the SILVA 138 16S rRNA database (Quast *et al*., 2013). Chloroplast and mitochondrial ASVs were filtered out from the datasets. Samples were rarefied at 45,000 reads before further analysis. Alpha diversity metrics, including Shannon diversity, Evenness, and Faith’s Phylogenetic Diversity (PD), were calculated to evaluate species richness and PD within each sample. Beta diversity was assessed by calculating pairwise community dissimilarity using the Bray-Curtis distance, followed by dimensionality reduction via principal coordinates analysis (PCoA) to visualise differences in microbial community structure between samples. Canonical correspondence analysis (CCA) was conducted using PAST (version 4.3) to explore the relationships between microbial community composition, functions, and environmental variables. Statistical differences between groups were evaluated using the Wilcoxon rank-sum test for two-group comparisons and the Kruskal-Wallis test for comparisons involving more than two groups, performed in R (version 4.2.3).

### 2.5 Functional prediction of microbial communities in mangrove sediments

To evaluate the predicted functions of the microbiome in mangrove soils, functional predictions were conducted using the PICRUSt2 pipeline (Douglas *et al*., 2020). This tool uses phylogenetic information to infer gene content and functional capabilities from microbial communities based on ASVs. ASV sequences were first aligned with reference genomes to predict the presence of functional genes. The predictions were normalised for 16S rRNA gene copy numbers using the castor R package (Louca & Doebeli, 2018) to reduce biases in functional gene prediction, which could result from varying gene copy numbers across taxa. ASVs with a nearest-sequence taxon index (NSTI) value above 2 were excluded from further analysis due to suspected spurious functional predictions. The functional predictions were then assigned to known metabolic pathways and gene families using the Kyoto Encyclopedia of Genes and Genomes (KEGG) database (Kanehisa *et al*., 2023), focusing on pathways related to carbon fixation, methane metabolism, and nitrogen and sulphur metabolism. Principle component analysis (PCA) was performed on predicted KEGG ortholog abundances to visualise functional variation among sediment samples.

### 2.6 Linking microbial taxa to functional abundance

To examine the relationships between microbial taxa and their predicted functional roles, functional profiles generated from PICRUSt2 were linked to the corresponding bacterial taxa based on KEGG Orthology (KO) annotations. Functional abundances related to carbon metabolism pathways were extracted and aggregated at the phylum level. To visualise the distribution of major microbial contributors across functions, a bubble plot was created using the ggplot2 package in R (version 4.2.3), where bubble size represented the relative abundance of each functional category contributed by different phyla.

### 2.7 Co-occurrence network analyses

Co-occurrence network analyses were performed to explore interactions among microbial taxa in mangrove soils. Microbial association networks were inferred using the SparCC function in the SpiecEasi package (version 1.13) in R (version 4.2.3), which accounts for compositionality and sparsity in microbial data. An adjacency matrix was constructed with a correlation threshold of 0.3 to retain significant associations. Networks were visualized using Gephi (version 10), where positive and negative correlations were represented by blue and red edges, respectively. Node sizes were scaled according to betweenness centrality scores, highlighting key taxa with potential ecological importance.

## 3. Results

### 3.1 Distinct physicochemical and biological properties across mangrove locations and depths

Physicochemical (temperature, pH, salinity, TOC, TN, sediment type) and biological (microbial cell abundance) properties from the mangrove sediment samples (SB, PL, KS) were collected (Table S1). The average mangrove soil temperature, pH, and salinity were 28-29 °C, 5.7, and 3.4 PSU, respectively. Salinity varies across sites (p<0.05), with KS exhibiting higher salinity at average 3.98 PSU compared to PL at 3.14 PSU and SB, at 2.59 PSU. Temperature also varied significantly (p<0.05) across sites where KS possessed the highest average temperature at 30 °C, whereas the lowest temperature was observed in SB at an average of 28 °C. Across depths, pH was found to be significantly higher at the top (average pH 6.6) than at the bottom (average pH 5.1) sediment (p<0.05). TOC, TN and cell abundance were significantly higher in PL compared to SB and KS. Depending on the particle size, the sediment samples were differentiated into three types of sediment textures: sand, silt and clay, with sand being the coarsest and clay being the finest. KS were dominantly silt-clay (50% silt, 40% clay), while SB had a greater proportion of sand (45%) and PL higher clay content (80%). The above data shows clear differences in most physicochemical and biological properties across the three mangrove locations (SB, PL, KS) and depths, suggesting distinct environmental conditions of each mangrove ecosystem.

### 3.2 Alpha and beta diversity of microbial communities in mangrove sediments

Based on the 16S rRNA amplicon sequencing data, a total of 6,089,044 reads were obtained, with an average of 190,283 reads per sample (Table S2). We rarefied the sequence at 45,000 sample reads as the curve of the bacterial 16S gene began to plateau (Fig. S1). The sequence data encompassed 69,569 Amplicon Sequence Variants (ASV), with an average of 2174 ASVs per sample. The alpha diversity indices revealed variations across different mangrove locations (Table S3). The Shannon diversity index was significantly higher (p <0.05) in SB (10.3) compared to PL (10.1) and KS (9.8). However, no significant differences were observed in the richness (number of ASVs), evenness, and Faith’s Phylogenetic Diversity indices across the mangrove locations and depths. These results indicate the microbial diversity across these sediment samples were largely consistent, although SB may harbour richer and more evenly distributed bacterial diversity compared to other locations. The beta diversity of microbial communities across the three mangrove sites (SB, PL, KS) based on Bray-Curtis’s dissimilarity matrix of the obtained ASVs showed distinct clustering of community composition based on locations (Fig. 3). This trend also applied to seagrass sediments (MB), which also had unique community assemblages from that of the mangrove sediments. Mock community samples (ZymoBIOMICS) that were sequenced along with the sediment samples in two sequencing runs also showed unique and consistent clustering separate from the environmental samples (Fig. S1).

Beyond location-based clustering, depth also contributed to microbial community segregation, although to a lesser extent. Samples from top, middle and bottom sediment layers showed partial separation within each site cluster (Fig. 3B). Top sediments generally grouped more closely together, while middle and bottom layers exhibited greater dispersion, particularly in KS and PL, suggesting increasing heterogeneity with sediment depth. These findings suggest that location acts as the primary driver of microbial community structure in mangrove sediment, while depth provides secondary but meaningful structuring, likely reflecting vertical physicochemical gradients within the sediment profile.

### 3.3 Taxonomic composition of mangrove sediment microbial communities and their relationship with environmental parameters

Based on the ASVs of the 16S rRNA gene sequences obtained, the microbial taxa present in all mangrove sediments consisted of 83 phyla, 230 classes and 550 orders. Overall, the ten most abundant phyla include Pseudomonadota (19.6%), Chloroflexota (19.59%), Desulfobacterota (14.2%), Acidobacteriota (7.5%), Actinobacteriota (6.97%), Planctomycetota (4.17%), Bacteroidota (3.38%), Gemmatimonadota (2.30%), Bacillota (2.21%), and Myxococcota (1.70%) (Figs. 3A and S3).

Although these phyla are present in all mangrove locations, significant differences in relative abundances across depths were evident for certain phyla, e.g., Pseudomonadota and Chloroflexota (Fig. S3) (P<0.05). Additionally, Planctomycetota was most prominent in KS compared to SB and PL, which had significantly lower abundance (Fig. S4) (P<0.05). These results indicate that whilst mangrove sediments consist of a core microbiome, some communities differ in abundance across different depths and locations, which contributes to the unique community composition observed (Fig. 3B). Our Canonical Correspondence Analysis (CCA) showed different environmental factors may drive these compositional differences (Fig. S6). Specifically, microbial communities in KS were strongly influenced by pH, clay % and conductivity, whereas those from SB and PL were more associated with particle size, sand % and cell abundance.

### 3.4 Predicted microbial functional diversity and nutrient metabolism in mangrove sediments

We analysed the 16S rRNA sequence using the PICRUSt functional prediction pipeline and identified a total of 407 modules of KEGG Orthology groups (KOs) in the mangrove sediment samples. Based on Bray-Curtis’s dissimilarity matrix of the KOs, the functional diversity of the mangrove sediments was shown to be largely clustered based on depth rather than locations of the mangrove sediments (Fig. 5A). CCA of the functional modules and environmental variables also highlighted this clustering, with different degrees of influence by pH, sediment type, depth and salinity potentially driving the functional composition across depths (Fig. S7).

Based on the predicted functional data, we also measured the abundance of key metabolic pathways (Fig. 5B), focusing on carbon and nutrient-related metabolism among the three mangrove locations. Our data revealed that carbon fixation exhibited the highest gene abundance in carbon metabolism, followed by carbohydrate metabolism, including glycolysis/gluconeogenesis, the citrate cycle, and methane metabolism. Meanwhile, sulfur and nitrogen metabolism exhibited the lowest gene abundance. Except for methane metabolism in SB, all predicted pathways were in similar abundance across the locations.

Genes contributing to specific microbial carbon fixation or assimilation pathways were abundant at varying depths of mangrove sediments (Fig. S8). In particular, the reductive pentose phosphate cycle (Calvin-Benson cycle) and 3-Hydroxypropionate cycles were prominent in the top layer, whilst the Wood-Ljungdahl cycle and methanogenesis were dominant in the bottom layer of the mangrove sediments.

### 3.4 Microbial group associated with carbon and nutrient metabolism in mangrove sediments

We further analysed the abundance of taxa most associated with several predicted carbon and nutrient-related metabolic pathways. Desulfobacterota emerged as the dominant contributor to nutrient metabolic processes, consistently exhibiting the highest involvement in carbon fixation, as well as methane, carbohydrate, sulfur, and nitrogen metabolism. Chloroflexota predominantly contributed to carbon fixation and methane metabolism, whereas Pseudomonadota showed high abundance in sulfur and nitrogen metabolism but contributed relatively to a lesser extent in carbon fixation and methane metabolism. Planctomycetota showed involvement in all four metabolic processes, especially in nitrogen metabolism. Crenarchaeota and archaeal phylum, showed a relatively higher involvement in methane metabolism.

### 3.5 Co-occurrence network analysis of microbial communities in mangrove sediments

Based on the sequences, we performed a co-occurrence network analysis, which elucidates possible interactions among microbial communities in mangrove sediments (Fig. S9). The network showed a high density of positive and negative connections, reflecting the complex interactions among the microorganisms. Pseudomonadota (Proteobacteria), Latescibacterota, Desulfobacterota and Chloroflexota (Chloroflexi), Planctomycetota and Actinobacterota emerged as among the keystone bacteria in the sediments, with the first four groups being the most prominent. An unclassified archaeal group showed most prominence among archaea, followed by Asgardarchaeota and Crenarchaeota.

## 4 Discussion

### 4.1 Mangrove sediments across different locations possess distinct environmental profiles and microbial community composition

Mangrove ecosystems are shaped by highly variable environmental conditions driven by tidal dynamics, freshwater input, and geomorphology (Alongi, 2002; Kristensen *et al*., 2008). Consistent with this, the three mangrove locations examined in this study, i.e., Seri Buat (SB), Sg Pulai (PL), and Kuala Selangor (KS), exhibited distinct physicochemical characteristics, including salinity, pH, sediment texture, organic carbon (OC), and total nitrogen (TN) (Fig. 2, Table S1), that may contribute to the unique microbial community composition observed across the three locations (Fig. 3B). Indeed, we observed that key environmental gradients, such as sediment type, pH and cell abundance, potentially drive the community composition to a different extent in these mangrove locations (Fig. S6). Such associations align with previous work demonstrating certain parameters (e.g., soil texture, pH, and nutrient availability) shape mangrove microbial assemblages, which support the idea that environmental heterogeneity is a dominant force shaping communities in mangrove sediments (Chen *et al*., 2016; Chambers *et al*., 2016).

**Fig 2.**
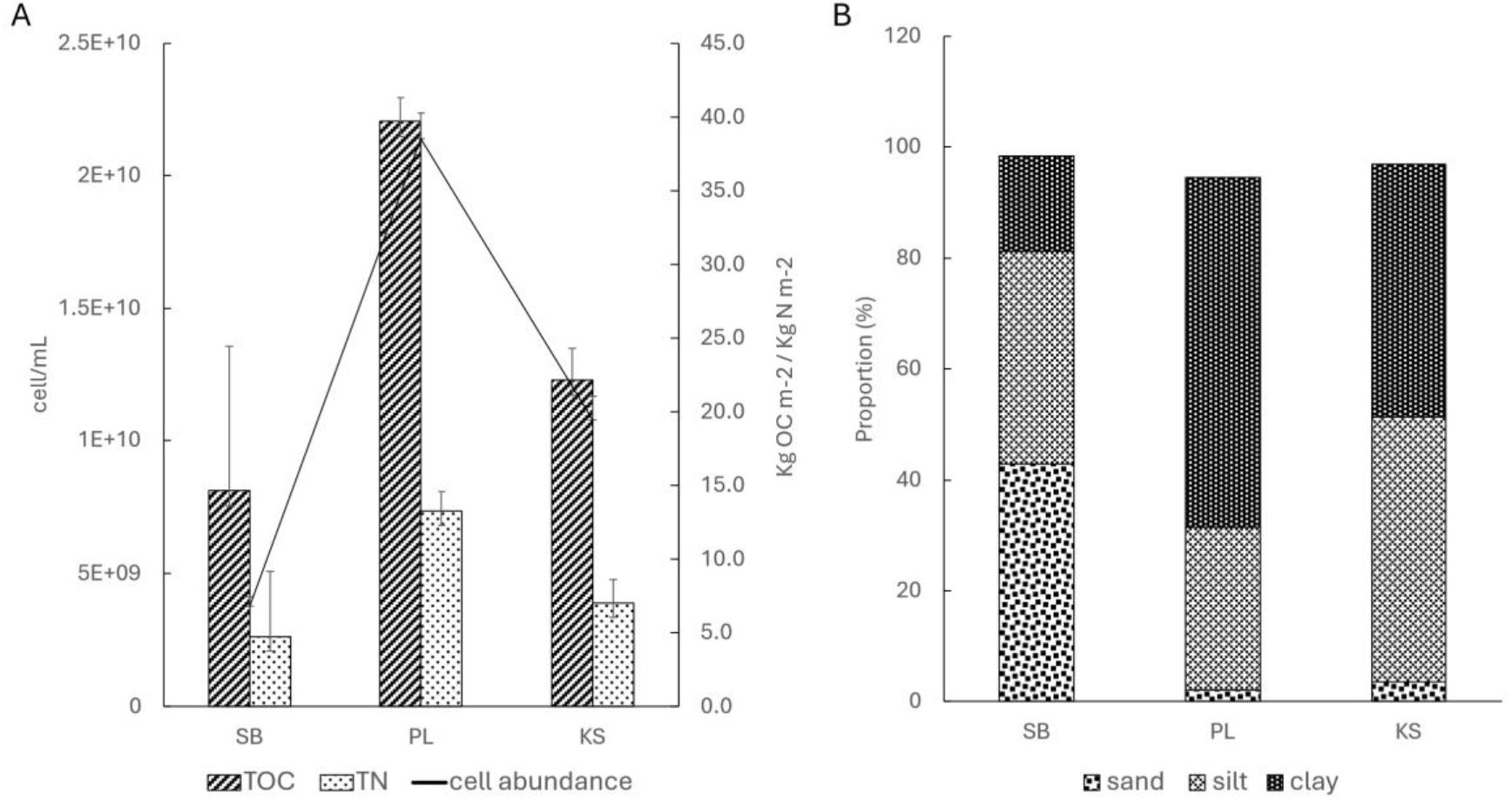
Physicochemical and biological parameters covering a) Total organic carbon (TOC), Total nitrogen (TN), and cell abundance (cells/mL) and b) sediment type proportion (%) across Seribuat Island (SB), Sg Pulai Estuary (PL), and Kuala Selangor (KS) mangrove sediments.

**Fig 3.**
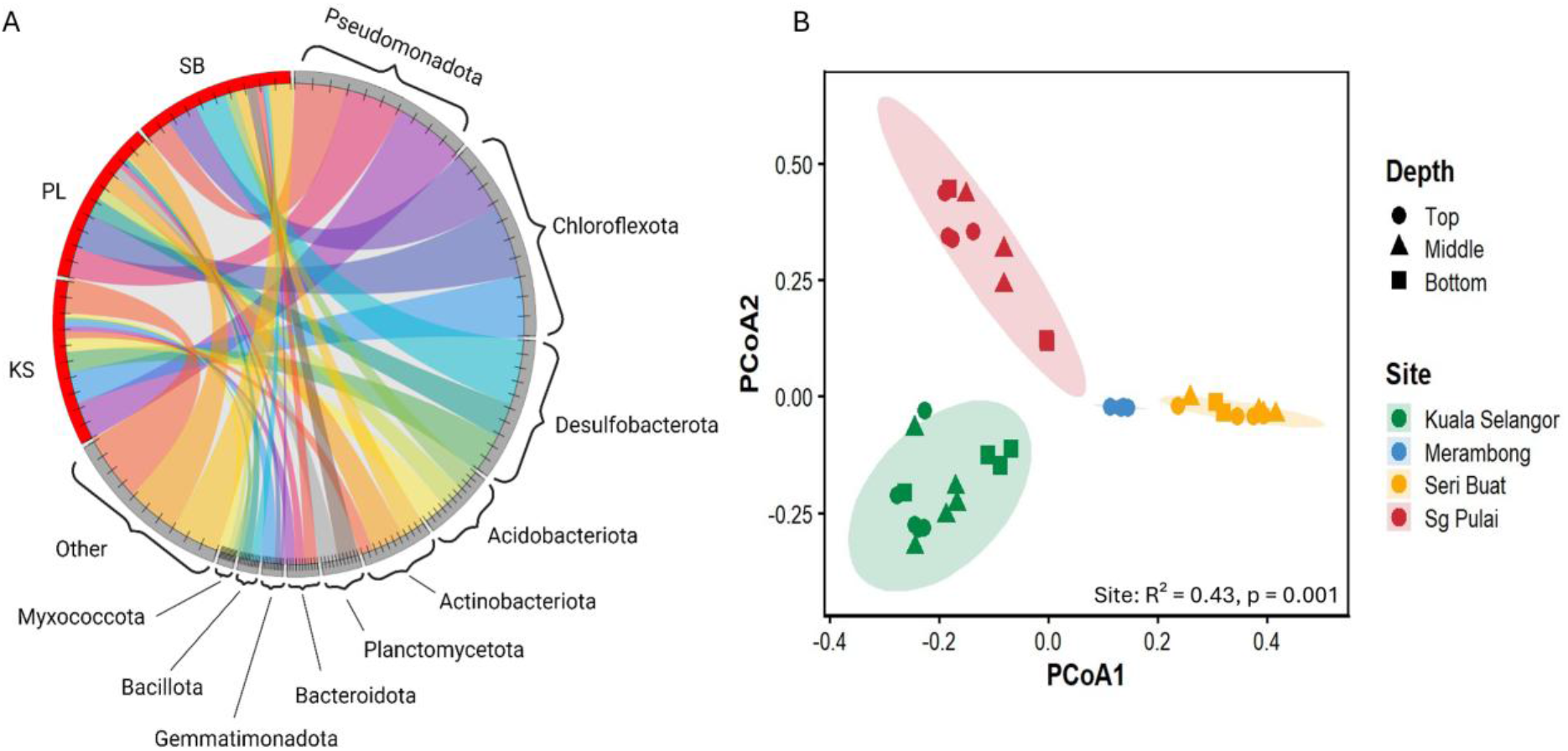
(A) Chord Plot of Top 10 most abundant: *Pseudomonadota, Chloroflexota, Desulfobacterota, Acidobacteriota, Actinobacteriota, Planctomycetota, Bacteroidota, Gemmatimonadota, Bacillota, and Myxococcota*, (B) A principal coordinates analysis (PCoA) plot based on the Bray-Curtis distance between each sample of microbial community composition across different mangrove sediments and depths; Yellow: Pulau Seri Buat (SB), Red: Sg Pulai (PL), Green: Kuala Selangor (KS).

In addition to physicochemical factors, mangrove vegetation may also influence sediment properties and microbial community composition. The dominant mangrove species differed across the study locations, with PL primarily dominated by *Rhizophora apiculata*, SB characterised by *Rhizophora apiculata* within a mixed *Rhizophora-Bruguiera* forest and KS dominated by *Brugueira parviflora*. Such variation in vegetation composition may influence sediment properties through differences in litter input, root architecture, and organic matter quality. Previous studies have demonstrated that mangrove species-specific traits, including root morphology and litter chemistry, significantly affect sediment biogeochemistry, nutrient availability, and microbial activity (Alongi, 2002; Kristensen *et al*., 2008; Reef *et al*., 2010). Collectively, this suggests that vegetation structure, alongside physicochemical gradients, plays an important role in shaping microbial assemblages and composition within mangrove sediments.

### 4.2 The mangrove sediment harbours a core microbiome linked to conserved biogeochemical functions

Despite clear spatial separation in microbial community composition among the three mangrove locations (Fig. 3B), a consistent core microbiome was observed across all locations (Fig. 3A), suggesting that mangrove sediments may maintain a conserved microbial assemblage that supports fundamental ecosystem functions. Shared dominant phyla across locations include Pseudomonadota, Desulfobacterota, Chloroflexota, Acidobacteriota, Actinobacteriota and Planctomycetota. These dominant taxa are consistent with previous studies reporting similar key microbial groups in Malaysian mangrove sediments (Aqeela *et al*., 2025). The persistence of these taxa across environmentally distinct mangrove systems suggests that, although environmental filtering shapes site-specific communities, a subset of microbial groups is selectively maintained due to their functional importance in mangrove sediment biogeochemistry.

Among shared taxa, Desulfobacterota were consistently abundant across all locations, highlighting their potential role as key mediators of anaerobic carbon remineralisation in mangrove sediments. Mangrove ecosystems are typically characterised by waterlogged and oxygen-limited conditions, which favour sulfate-reducing microorganisms that utilise sulfate as an alternative electron acceptor during organic matter degradation. The dominance of Desulfobacterota observed in this study aligns with previous reports demonstrating that sulfate-reducing bacteria represent one of the most important microbial functional groups in mangrove sediments, contributing to organic carbon turnover and influencing carbon preservation under anoxic conditions (Wu *et al*., 2019; Li *et al*., 2021; Wu *et al*., 2021).

Similarly, Chloroflexota were also identified as part of the shared microbiomes across study locations. Their presence with varying organic carbon levels suggests that Chloroflexota may contribute to the breakdown of complex plant-derived organic matter, including lignocellulosic compounds. Previous studies have reported Chloroflexota as persistent members of mangrove sediment microbiomes, often linked to slow organic matter degradation and syntrophic interactions with other anaerobic microorganisms (Hug *et al*., 2013; Wasmund *et al*., 2016; Zhou *et al*., 2017). The coexistence of Chloroflexota with sulfate-reducing Desulfobacterota in this study further suggests potential metabolic coupling where fermentation products generated by Chloflexota may serve as substrates for sulfate-reducing bacteria, thereby supporting interconnected carbon cycling pathways in mangrove sediments.

The persistence of Pseudomonadota across environmentally distinct mangrove sediments suggests their ecological versatility and ability to fluctuating redox conditions driven by tidal cycles. Their widespread occurrence in this study supports the notion that metabolically flexible microbial groups may form an essential component of the core microbiome, enabling mangrove ecosystems to maintain functional stability under changing environmental conditions.

The presence of these shared microbial taxa across environmentally distinct mangrove locations suggests that mangrove sediments may harbour a functionally conserved microbiome that supports essential biogeochemical processes particularly carbon and nutrient cycling. While environmental heterogeneity shapes community composition at each location, the persistence of these core microbial groups indicates potential functional redundancy within mangrove ecosystems. This observation is consistent with previous studies demonstrating that mangrove microbiomes exhibit both compositional variability and functional stability, highlighting the importance of core microbial communities in sustaining mangrove ecosystem functioning (Chen *et al*., 2016; Wu *et al*., 2021).

### 4.3 Microbial functional diversity in mangrove sediments remain conserved across locations but varied across depths

Consistent with the presence of a shared core microbiome across the three mangrove locations, predicted functional analysis showed that the functional composition in these locations remains conserved, only differing more prominently based on depth (Fig. 5A). This suggests that while taxonomic composition varies spatially, microbial communities in mangrove sediments may exhibit functional redundancy, whereby different taxa perform similar ecological roles to maintain ecosystem functioning. Such functional conservation has been widely reported in mangrove ecosystems, where microbial communities maintain stable biogeochemical processes despite environmental heterogeneity (Louca *et al*., 2018; Chen *et al*., 2016).

**Fig 5.**
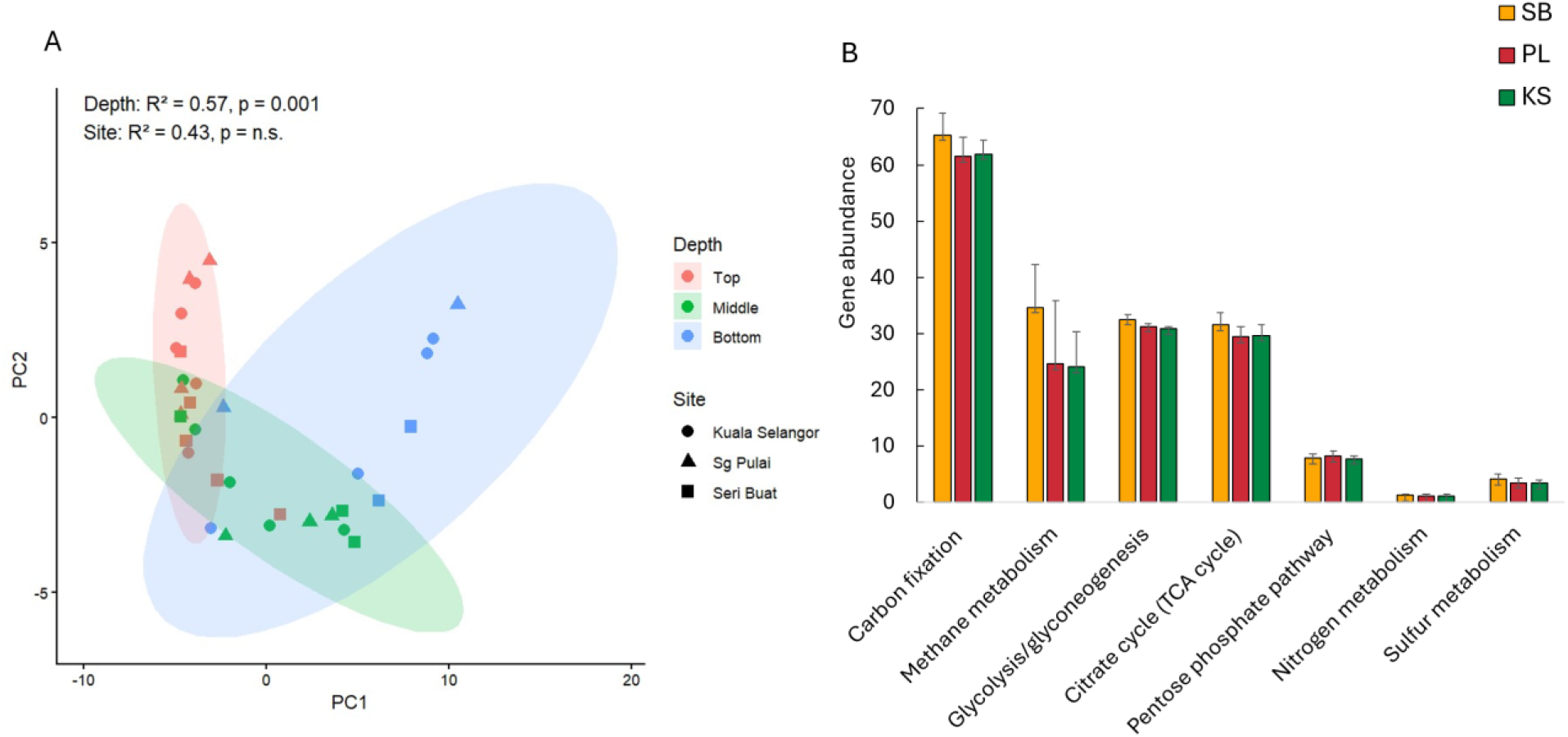
A) Principal Component Analysis (PCA) of predicted microbial functional composition across different depths and locations (KS, PL, SB) and depths (Top, Middle, Bottom). B) Predicted gene abundance on carbon and nutrient-related metabolism across three different mangrove sediments.

Across all locations, dominant predicted functions included carbon fixation, methane metabolism, glycolysis/gluconeogenesis, the tricarboxylic acid (TCA) cycle, nitrogen metabolism, phosphate metabolism and sulfur metabolism (Fig. 5B). These pathways collectively represent fundamental metabolic processes supporting carbon turnover, nutrient cycling, and energy production in mangrove sediments. The relatively similar abundance of these pathways across the three locations suggests that core ecosystem functions are maintained regardless of location-specific environmental conditions. This observation aligns with previous studies demonstrating that mangrove sediments harbour metabolically versatile microbial communities capable of sustaining key biogeochemical cycles (Alongi, 2014; Wang *et al*., 2025).

In contrast, depth emerged as a stronger determinant of functional distribution. Vertical gradients in mangrove sediments typically influence oxygen availability, redox potential, sulphate concentration, and organic matter composition, all of which regulate microbial metabolic processes (Kristensen *et al*., 2008; Alongi, 2014). These depth-related environmental gradients were also reflected in our predicted functional profiles, suggesting functional stratification of microbial communities along sediment depth.

Carbon fixation pathways displayed clear depth partitioning. Predicted gene abundance for the Calvin–Benson and 3-hydroxypropionate cycles were more abundant in surface sediments, where light penetration, fresher organic inputs, and higher oxygen availability favour phototrophic and aerobic autotrophic processes (Fig. S8). In contrast, deeper sediment layers were enriched in pathways characteristic of anaerobic carbon fixation, especially the Wood–Ljungdahl (WL) pathway (Fig. S8), which is typical of anaerobic bacteria and archaea capable of acetogenesis and hydrogenotrophy (Jiang *et al*., 2022). Methanogenesis predicted genes were also more concentrated in deeper sediments, while in contrast, genes related to methane oxidation were more abundant at the surface layers (Fig. S8). These results indicate the depth-specific functional zonation of the microbial communities, which, if disrupted, would potentially alter carbon sequestration and storage capacities of mangrove sediments and affect mangroves’ role as a vital carbon sink (Macreadie *et al*., 2025).

### 4.4 Prominent carbon-related pathways in the mangrove sediment

As mangrove sediments conserved their microbial function, we further explored the gene abundance related to carbon metabolism. This study mainly focuses on several key carbon metabolic pathways such as carbon fixation, glycolysis/gluconeogenesis, and the TCA cycle. Previous study showed that enhanced microbial diversity would promote the transformation and utilization of OC (Berendsen *et al*., 2012). The findings of this study showed a similar approach where the highest bacterial diversity in SB and PL exhibited the higher gene abundances involved in carbon-related pathways. A current study revealed that carbon fixation exhibits the highest gene abundance across carbon-related pathways. This carbon fixation is primarily mediated by autotrophic microorganisms which play a crucial role in the mangrove ecosystems by converting inorganic carbon into organic compounds. This process is vital for long-term carbon sequestration, as it ensures that more carbon is stored in the sediment than released in the atmosphere.

In this study, we observed that carbon fixation dominated over carbon decomposition (Fig 5). Carbon metabolism is a key driver of carbon decomposition in mangrove sediment which the heterotrophic microorganisms facilitate the degradation of this carbohydrate by releasing CO_2_ via mineralization or incorporating the product into microbial biomass. This balance is essential for maintaining long-term carbon sequestration, as it prevents the release of stored carbon back into the atmosphere. If decomposition were to surpass fixation, carbon reservoirs could become destabilized, leading to the release of substantial amounts of CO_2_ and CH_4_, which would contribute to global warming. In addition, methane metabolism is part of carbon sequestration as the process of methanogenesis is the production of methane (CH_4_) which typically occurs in anoxic conditions. While methanogenesis contributes to greenhouse gas emissions, its impact is mitigated by methane-oxidizing bacteria, which consume CH_4_ before it can escape into the atmosphere. The current findings found that each depth has a significant abundance of methanogenesis and methane oxidation at the bottom and top layers respectively. This indicates that the redox potential of CH_4_ emission to the atmosphere is low, which is mostly due to large sulfate input, which promotes sulfate reduction and inhibits CH_4_ generation (Callaway *et al*., 2012). The prominence of carbon fixation in mangrove sediments has profound implications for environmental stability and climate regulation. Mangroves are crucial ecosystems in mitigating climate change by acting as long-term carbon reservoirs, buffering the rise in atmospheric CO_2_ levels. Moreover, the suppression of methane emissions through methane oxidation further reduces the greenhouse gas footprint of mangrove systems.

### 4.5 Desulfobacterota and Chloroflexota as dominant microbial drivers of carbon cycling in mangrove sediments

Across sites and depths, several microbial groups consistently emerged as major contributors to carbon-related pathways. Desulfobacterota is found to be the most prominent group, contributing heavily to carbon fixation, methane metabolism, and sulfur and nitrogen cycling (Fig. 6). Although best known as sulfate-reducing bacteria (SRB), many species and genera within phylum Desulfobacterota also possess genes (often through the WL pathway) for autotrophic carbon fixation, which enables them to assimilate inorganic carbon under anoxic, sulfate-rich conditions (Meyer & Kuever, 2007; Zhu *et al*., 2018). Their dual role in organic matter degradation and autotrophic carbon assimilation positions them as central players in carbon sequestration in anoxic mangrove sediments.

**Fig 6.**
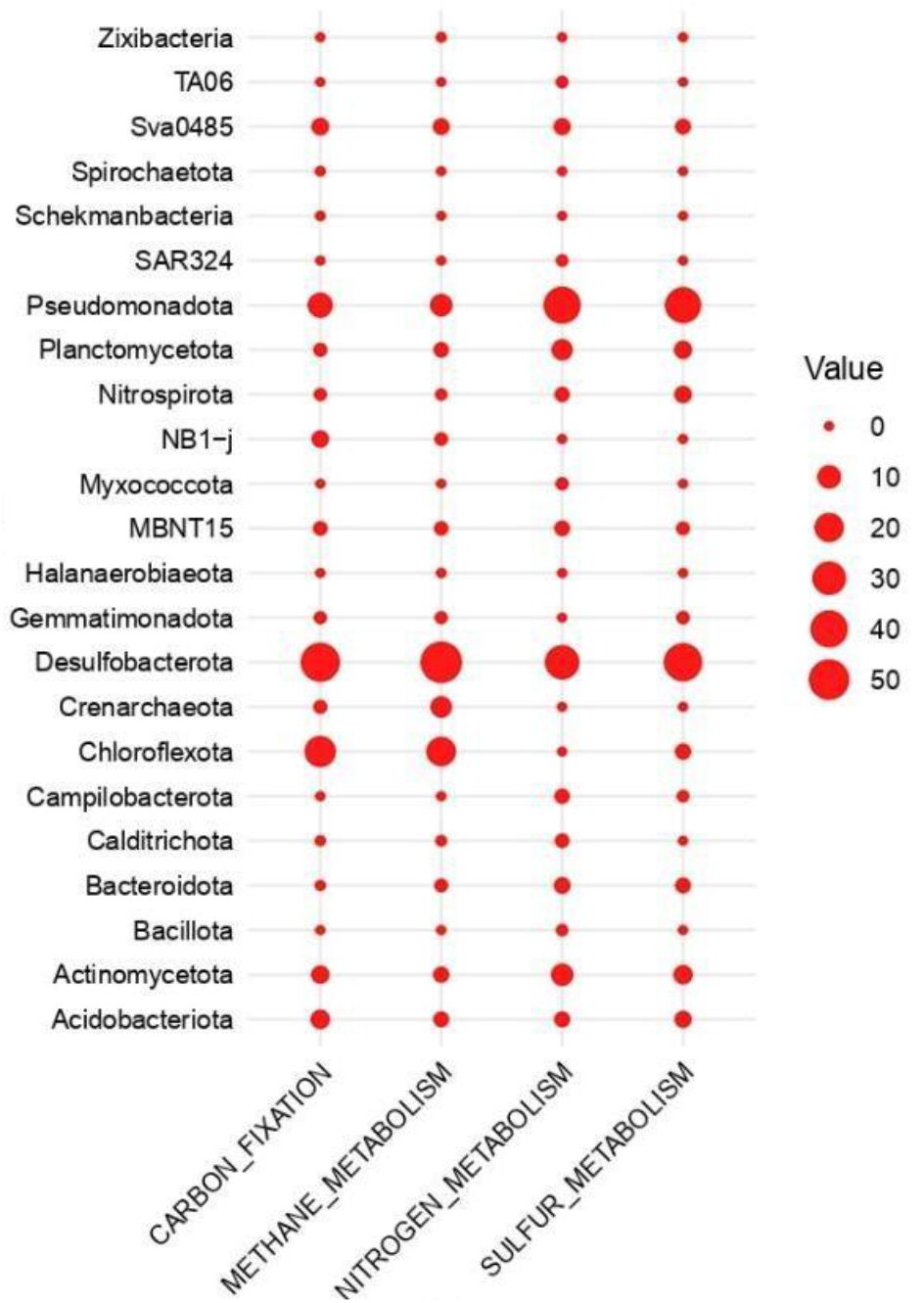
Contribution analysis of dominant microbial taxa to the function at the phylum level.

Chloroflexota also contributes substantially to the carbon cycle, mainly through carbon fixation and methane metabolism. Members of this phylum include both phototrophic and non-phototrophic lineages capable of operating under low-oxygen or anoxic conditions, frequently using the reductive TCA (rTCA) cycle for autotrophy (Jiang *et al*., 2022). Their enrichment in deeper sediments suggests a role in sustaining carbon assimilation under conditions where oxygen-dependent pathways are restricted. Co-occurrence network analysis further highlighted Desulfobacterota, Chloroflexota, Pseudomonadota, and Planctomycetota as keystone taxa (Fig. 7), suggesting the strong relationship between metabolically diverse microbial groups that promote the stability of sediment carbon processes. Although bacteria dominated the dataset, archaea, particularly Crenarchaeota, were predicted to contribute significantly to methane metabolism. Many crenarchaeal lineages participate in methanogenesis or anaerobic methane oxidation, processes central to determining greenhouse gas emissions from coastal wetlands (Holguin *et al*., 2001). Their detection suggests that archaeal communities, although less abundant, may exert disproportionate control over methane dynamics, highlighting the need for archaeal-inclusive analyses to better quantify their functional contributions to carbon cycling and greenhouse gas regulation in mangrove ecosystems.

## 5. Broader Implications and Future Research

The findings from this study have broad implications for understanding the role of microbial communities in mangrove soils and their contribution to global carbon cycling. One of the broader implications is the ecosystem-level influences of microbial communities on mangrove soil function. Microbes play a critical role in maintaining the stability and resilience of these ecosystems, particularly through their involvement in carbon cycling. By fixing inorganic carbon, metabolising organic matter, and interacting with various geochemical cycles (such as sulfur and nitrogen cycles), microbial communities ensure the sustained carbon storage capacity of mangrove soils. Furthermore, the activities of these microbes contribute to ecosystem services such as nutrient recycling and sediment stabilisation, which are crucial for the health and longevity of mangrove forests.

This research also highlights the interconnectedness of microbial taxa within the mangrove ecosystem. The interactions between autotrophic and heterotrophic microorganisms, as well as the contributions of both bacteria and archaea, emphasise the complexity of microbial networks that govern carbon cycling in these environments. These findings suggest that a holistic approach, which considers the functional diversity of the microbial community, is necessary for understanding the overall contribution of mangrove ecosystems to carbon sequestration.

Despite advances in our understanding of microbial processes in mangrove soils, significant knowledge gaps remain that require further exploration. One key area is the spatial and temporal dynamics of microbial communities in these ecosystems. Mangrove sediments are highly heterogeneous, with environmental factors such as salinity, oxygen availability, and organic matter content varying across depths and locations. Long-term studies are essential to assess how microbial communities and their functional roles evolve in response to seasonal changes, pollution, deforestation, and climate change. These studies would provide insights into the adaptive mechanisms that microbes employ under fluctuating environmental conditions, revealing their potential resilience or vulnerability to stress.

Another promising avenue involves multi-omics approaches to comprehensively investigate microbial functional potential. While current studies offer valuable information on the taxa involved in carbon fixation, integrating metagenomics, metatranscriptomics, metaproteomics, and metabolomics would allow for a real-time assessment of microbial activity and the metabolic pathways at play. This integrated approach could help link microbial diversity to specific functional outcomes, offering a deeper understanding of how different microbial taxa contribute to crucial processes such as carbon sequestration and nutrient cycling in mangrove ecosystems.

Finally, there is a need to develop predictive models that incorporate microbial dynamics into assessments of carbon sequestration potential in mangrove ecosystems. By integrating microbial data with environmental factors, hydrological processes, and climate projections, researchers can better predict how mangrove ecosystems will respond to future climate scenarios. Such models would be valuable tools for policymakers and conservationists, enabling more effective mangrove restoration and protection strategies, which are critical to maximising the role of these ecosystems as carbon sinks in a changing world.

## 5. Conclusion

This study provides key insights into the taxonomic and functional diversity of microbial communities in mangrove sediments, suggesting their role in carbon cycling across different environmental conditions. We found that despite significant compositional differences in microbial communities, their functional capabilities, particularly those related to carbon cycling, remain largely conserved. The strong influence of site-specific physicochemical factors such as salinity, organic carbon content, and soil texture on microbial community structure highlights the unique adaptability of these communities to local environmental conditions.

Our findings demonstrate the central role of Desulfobacterota and Chloroflexota in carbon metabolism, with both taxa contributing significantly to the ecological processes that underpin long-term carbon storage. The presence of functional redundancy further ensures the persistence of key biogeochemical functions, such as carbon fixation and organic matter decomposition, across diverse microbial communities. Together, these conserved functional traits and adaptive microbial responses emphasise the stability and consistency of microbial contributions to carbon cycling in mangrove ecosystems.

## Acknowledgements

We would like to thank all laboratory members and field assistants involved in sample collection and laboratory analyses.

