## Supplementary for "Microbial Diversity and Function Linked to Carbon Cycling in Mangrove Sediments"

**Table S1** Physicochemical metadata of mangrove sediment samples across different sites (SB, Seri Buat; PL, Sg Pulai; KS, Kuala Selangor) and depths (10 – 100 cm).

| Sample | Temperature (°C) | pH | Salinity (psu) | Conductivity  (mS/cm) | Depth (cm) | TN (Kg m-2) | OC (Kg m-2) | moisture | sand (%) | Silt  ( %) | clay (%) |
| --- | --- | --- | --- | --- | --- | --- | --- | --- | --- | --- | --- |
| SB-C1-L1 | 28.4 | 7.13 | 2.01 | 3833 | 10 | 2.45 | 49.76 | 44.06 | 75.9 | 12.63 | 11.35 |
| SB-C1-L3 | 28.6 | 6.99 | 1.97 | 3792 | 30 | 4.62 | 128.29 | 62.81 | 9.781 | 31.93 | 58.296 |
| SB-C1-L5 | 28.5 | 5.87 | 0.91 | 1784 | 50 | 4.27 | 164.19 | 68.8 | 16.86 | 79.95 | 3.2 |
| SB-C2-L1 | 28.1 | 6.08 | 1.32 | 5292 | 10 | 3.67 | 70.95 | 37.71 | 13.64 | 41.8 | 44.555 |
| SB-C2-L4 | 28.4 | 4.28 | 2.43 | 4625 | 40 | 3.86 | 122.87 | 56.04 | 15.68 | 49.02 | 35.3 |
| SB-C2-L8 | 28.1 | 4.87 | 2.21 | 4172 | 80 | 1.77 | 142.07 | 33.28 | 84.79 | 6.33 | 8.76 |
| SB-C3-L1 | 28.1 | 7.14 | 3.52 | 6450 | 10 | 4.32 | 111.57 | 60.44 | 77.93 | 21 | 1.07 |
| SB-C3-L5 | 28.1 | 6.55 | 5.02 | 9050 | 50 | 5.68 | 155.48 | 85.89 | 45.08 | 51.41 | 3.52 |
| SB-C3-L10 | 28.1 | 7.12 | 3.96 | 7199 | 100 | 4.89 | 116.72 | 67.06 | 46.11 | 52.01 | 1.89 |
| PL-C1-L1 | 29.3 | 5.66 | 1.98 | 3785 | 10 | 15.63 | 403.14 | 72.65 | 1.51 | 16.26 | 78.06 |
| PL-C1-L4 | 29.3 | 3.86 | 3.26 | 6043 | 40 | 16.13 | 509.51 | 98.56 | 1.12 | 52.86 | 41.94 |
| PL-C1-L7 | 29.3 | 3.83 | 4.34 | 7886 | 80 | 17.62 | 618.8 | 106.61 | 2.65 | 65.2 | 30.34 |
| PL-C2-L1 | 29.3 | 7.4 | 1.11 | 2158 | 10 | 8.28 | 149.48 | 52.3 | 1.44 | 0.14 | 98.42 |
| PL-C2-L4 | 28.6 | 3.53 | 2.53 | 4763 | 40 | 7.57 | 214 | 53.28 | 3.35 | 34.18 | 62.475 |
| PL-C2-L8 | 28.7 | 4.1 | 2.41 | 4462 | 80 | 11.09 | 354.98 | 57.1 | 0.85 | 5.31 | 87.8 |
| PL-C3-L1 | 29.3 | 7.15 | 1.03 | 2163 | 10 | 13.52 | 280.13 | 73.87 | 2.51 | 3.42 | 84.8 |
| PL-C3-L5 | 28.5 | 4.17 | 5.35 | 9654 | 50 | 10.39 | 470.26 | 105.56 | 2.17 | 12.68 | 76.95 |
| PL-C3-L10 | 28.7 | 3.6 | 6.3 | 11.2 | 100 | 18.34 | 483.76 | 99.02 | 3.09 | 74.42 | 21.66 |
| KS-C1-L1 | 29 | 6.07 | 4.29 | 7784 | 10 | 23.99 | 559.1 | 99.6 | 1.77 | 44.6 | 48.87 |
| KS-C1-L5 | 29 | 4.66 | 4.78 | 8765 | 50 | 13.3 | 333.03 | 79.28 | 8.79 | 71.52 | 19.31 |
| KS-C1-L10 | 29 | 4.43 | 4.38 | 7950 | 100 | 13.86 | 222.07 | 86.13 | 0.82 | 81.09 | 17.75 |
| KS-C2-L1 | 30 | 6.41 | 5.27 | 9506 | 10 | 16.36 | 199.93 | 91.32 | 2.62 | 32.19 | 60.74 |
| KS-C2-L5 | 30 | 4.32 | 4.11 | 7429 | 50 | 15.15 | 243.56 | 95.09 | 6.99 | 40.22 | 47.52 |
| KS-C2-L10 | 28 | 4.81 | 4.32 | 7838 | 100 | 13.23 | 158.76 | 102.24 | 4.4 | 59.53 | 33.74 |
| KS-C3-L1 | 30 | 6.25 | 2.79 | 5210 | 10 | 20.22 | 248.35 | 90.36 | 1.69 | 21.51 | 72.85 |
| KS-C3-L5 | 29 | 7.25 | 1.98 | 3763 | 50 | 14.21 | 177.72 | 83.46 | 1.56 | 28.64 | 65.13 |
| KS-C3-L10 | 31 | 5.45 | 3.27 | 6952 | 100 | 14.33 | 176.2 | 70.03 | 10.28 | 85.79 | 3.93 |
| KS-C4-L2 | 30 | 6.7 | 4.03 | 7526 | 10 | 16.16 | 213.97 | 80.5 | 1.55 | 5.64 | 86.6 |
| KS-C4-L5 | 30 | 6.77 | 3.73 | 7958 | 50 | 14.37 | 172.32 | 82.63 | 1.11 | 23.74 | 71.16 |
| KS-C5-L1 | 30 | 6.91 | 4.05 | 7762 | 10 | 18.08 | 190.12 | 91.73 | 4.76 | 62.73 | 31.08 |
| KS-C5-L5 | 30 | 7.05 | 4.77 | 7645 | 50 | 14.71 | 186.69 | 78.83 | 2.21 | 60.21 | 35.41 |
| KS-C5-L8 | 30 | 7.12 | 4.06 | 7869 | 80 | 14.94 | 175.04 | 75.37 | 1.5 | 51.31 | 45.45 |

**Table S2** Sequencing depth and rarefied read counts of mangrove sediment samples analysed.

| Sample | Raw Reads | Rarefied |
| --- | --- | --- |
| PL-C1-L1 | 206089 | 45000 |
| PL-C1-L4 | 192878 | 45000 |
| PL-C1-L7 | 113935 | 45000 |
| PL-C2-L1 | 136532 | 45000 |
| PL-C2-L4 | 255033 | 45000 |
| PL-C2-L8 | 288009 | 45000 |
| PL-C3-L1 | 181069 | 45000 |
| PL-C3-L10 | 195324 | 45000 |
| PL-C3-L5 | 221961 | 45000 |
| SB-C1-L1 | 134745 | 45000 |
| SB-C1-L4 | 119667 | 45000 |
| SB-C1-L8 | 115108 | 45000 |
| SB-C2-L1 | 128740 | 45000 |
| SB-C2-L4 | 168981 | 45000 |
| SB-C2-L8 | 181583 | 45000 |
| SB-C3-L1 | 113771 | 45000 |
| SB-C3-L10 | 250480 | 45000 |
| SB-C3-L5 | 258253 | 45000 |
| KS-C1-L10 | 201280 | 45000 |
| KS-C1-L1 | 202954 | 45000 |
| KS-C1-L5 | 200542 | 45000 |
| KS-C2-L10 | 203428 | 45000 |
| KS-C2-L1 | 200080 | 45000 |
| KS-C2-L5 | 201968 | 45000 |
| KS-C3-L10 | 203276 | 45000 |
| KS-C3-L1 | 203228 | 45000 |
| KS-C3-L5 | 202092 | 45000 |
| KS-C4-L2 | 201868 | 45000 |
| KS-C4-L5 | 203716 | 45000 |
| KS-C5-L1 | 200218 | 45000 |
| KS-C5-L5 | 200144 | 45000 |
| KS-C5-L8 | 202092 | 45000 |

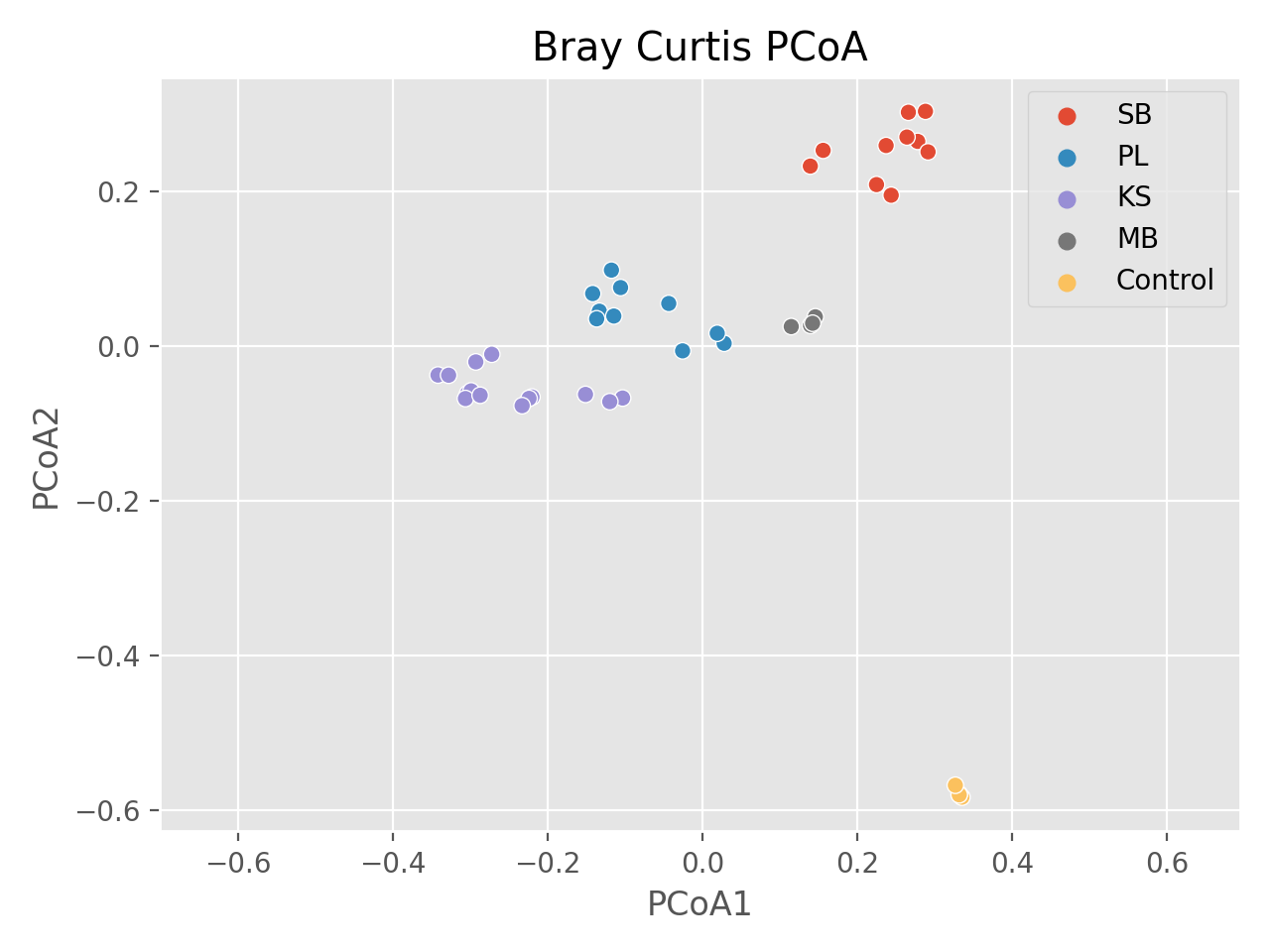

**Fig. S1** Bray-Curtis’s dissimilarity matrix from three different mangrove sites (SB, PL and KS), seagrass (MB) and control samples consisting of a known mock community (zymoBIOMICS).

**Table S3** Comparison of alpha diversity index of 16S rRNA sequencing from mangrove sediments (SB, PL, KS).

| **Sample** | **ASV** | **Shannon** | **Evenness** | **Faith PD** |
| --- | --- | --- | --- | --- |
| SB-C1-L1 | 2041 | 10.25918 | 0.933064 | 178.022 |
| SB-C1-L3 | 1427 | 9.704676 | 0.926133 | 133.7389 |
| SB-C1-L5 | 1189 | 9.494996 | 0.929522 | 119.5899 |
| SB-C2-L1 | 1140 | 9.425307 | 0.928146 | 112.5035 |
| SB-C2-L4 | 2160 | 10.3167 | 0.931399 | 188.3646 |
| SB-C2-L8 | 3076 | 10.78554 | 0.930835 | 243.1094 |
| SB-C3-L1 | 1592 | 9.844039 | 0.925546 | 152.6124 |
| SB-C3-L5 | 3992 | 11.17135 | 0.933895 | 262.2863 |
| SB-C3-L10 | 3766 | 11.00393 | 0.926394 | 251.4775 |
| PL-C1-L1 | 2300 | 10.16181 | 0.909994 | 176.3484 |
| PL-C1-L4 | 3208 | 10.60048 | 0.910156 | 251.9696 |
| PL-C1-L7 | 1353 | 9.253615 | 0.889663 | 135.1958 |
| PL-C2-L1 | 1926 | 10.12603 | 0.928043 | 149.8873 |
| PL-C2-L4 | 2164 | 9.203197 | 0.830731 | 175.5371 |
| PL-C2-L8 | 3025 | 10.25753 | 0.887191 | 224.6877 |
| PL-C3-L1 | 2189 | 10.18423 | 0.917856 | 162.6626 |
| PL-C3-L5 | 2734 | 9.916337 | 0.868637 | 198.6139 |
| PL-C3-L10 | 2295 | 9.917079 | 0.888353 | 180.5668 |
| KS-C1-L1 | 1953 | 9.865385 | 0.889689 | 159.5867 |
| KS-C1-L5 | 2009 | 9.405505 | 0.844398 | 166.6121 |
| KS-C1-L10 | 2183 | 10.06165 | 0.894823 | 172.7233 |
| KS-C2-L1 | 1952 | 9.788172 | 0.883365 | 157.6958 |
| KS-C2-L5 | 1696 | 9.209044 | 0.841104 | 158.5105 |
| KS-C2-L10 | 1872 | 9.32943 | 0.840831 | 160.7567 |
| KS-C3-L1 | 2383 | 10.11821 | 0.884534 | 193.1489 |
| KS-C3-L5 | 2385 | 10.18023 | 0.887894 | 192.1176 |
| KS-C3-L10 | 2233 | 9.810761 | 0.869666 | 172.4807 |
| KS-C4-L2 | 1999 | 9.831032 | 0.884427 | 155.7809 |
| KS-C4-L5 | 1796 | 9.538323 | 0.871223 | 153.4242 |
| KS-C5-L1 | 2007 | 9.805187 | 0.878579 | 160.7123 |
| KS-C5-L5 | 1697 | 9.278726 | 0.855846 | 150.2027 |
| KS-C5-L8 | 1827 | 9.359784 | 0.854488 | 150.4803 |

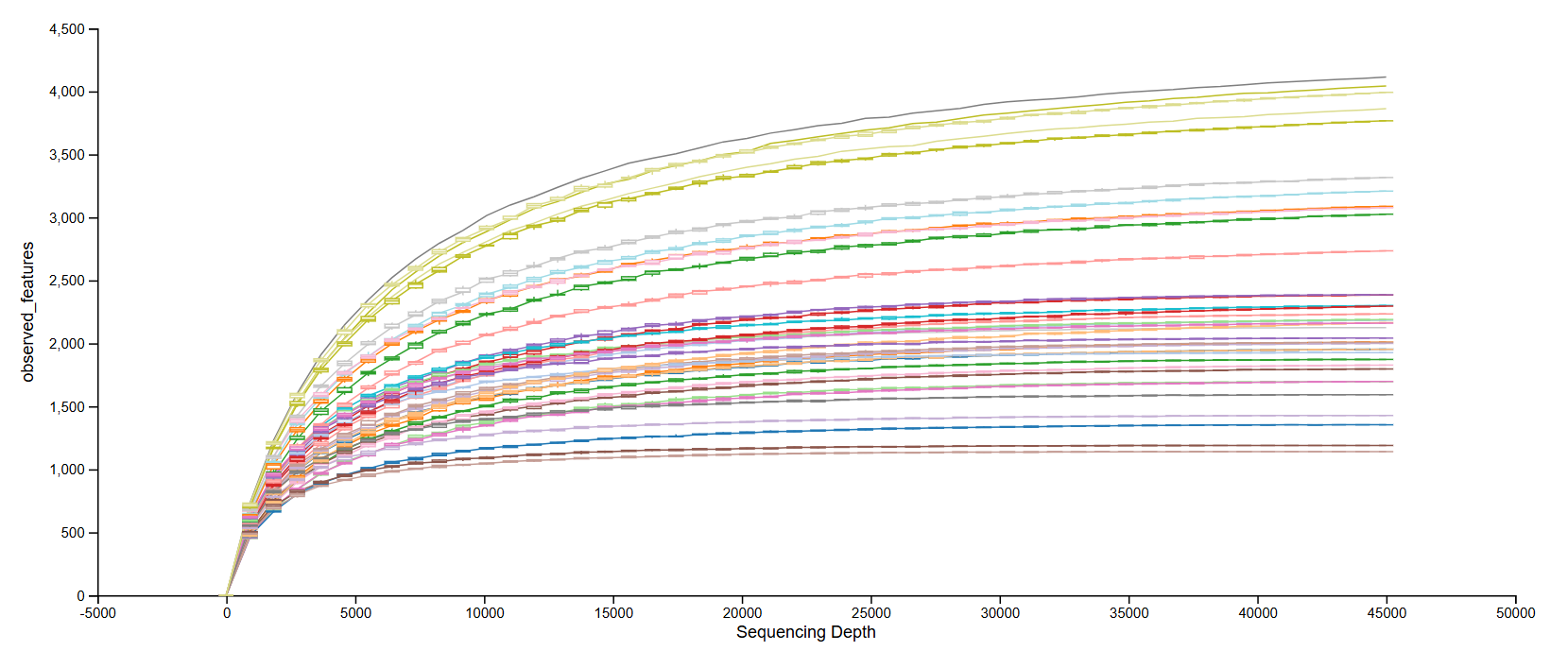

**Fig. S2** Rarefaction curves of all mangrove sediment samples from PL, SB, and KS, showing sequencing depth and observed microbial diversity across samples.

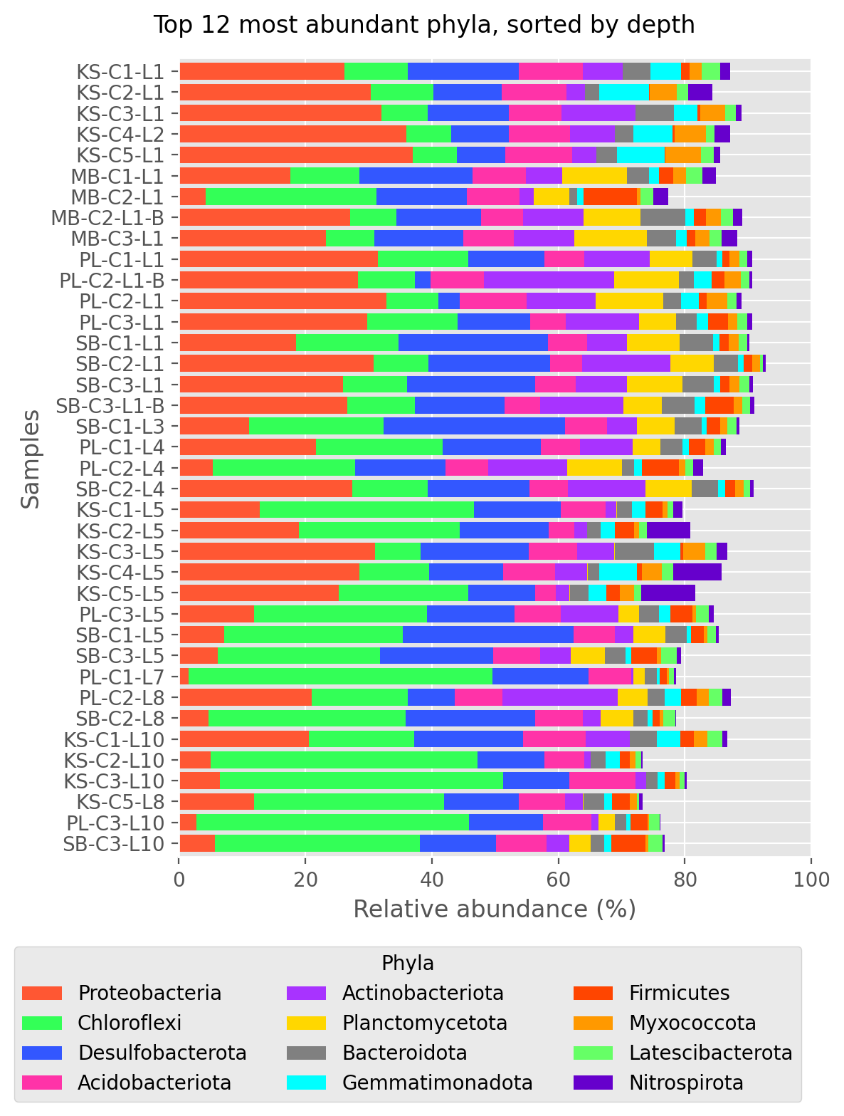

**Fig. S3** Top twelve most abundant phyla from all samples (SB, PL, KS)

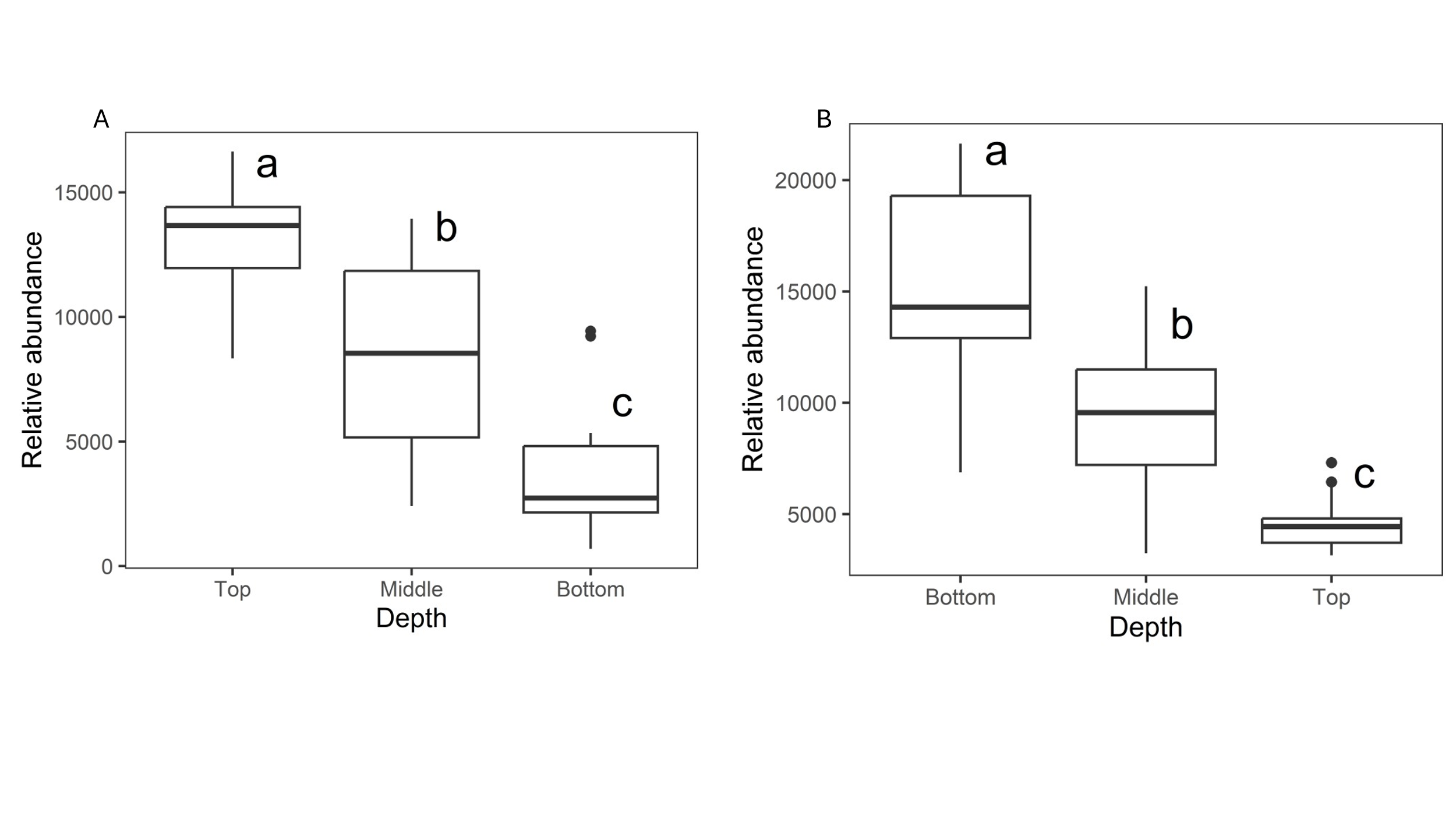
**Fig. S4** One-way ANOVA analysis showing significant differences in the relative abundance of phyla (A) Pseudomonadota and (B) Chloroflexota across sediment depths (bottom, middle, top).

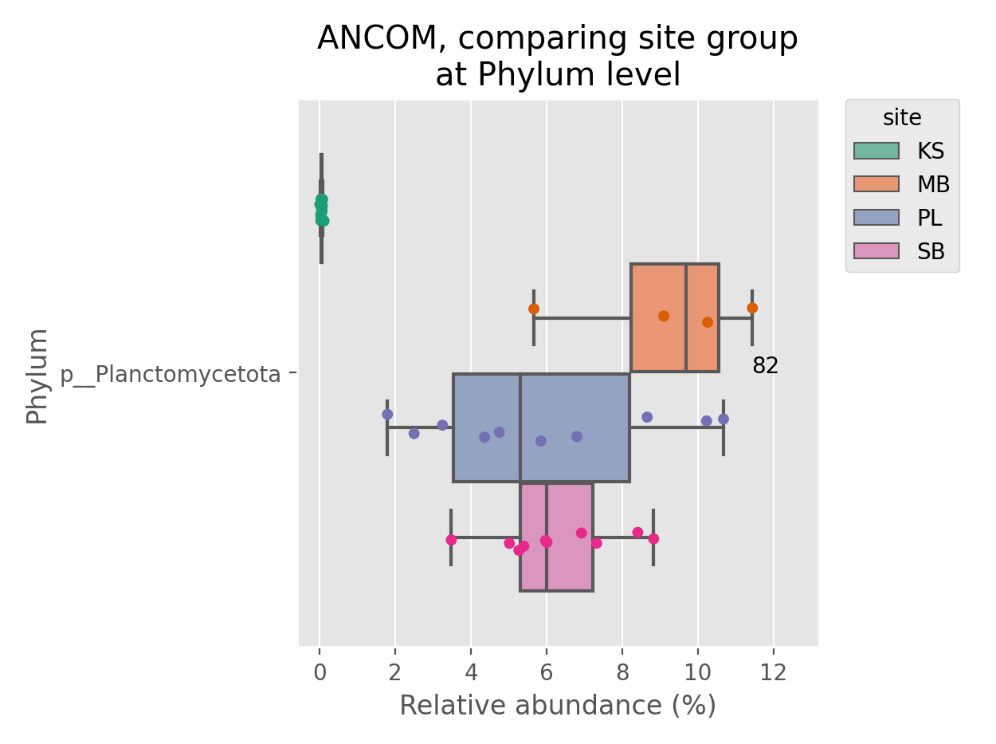

**Fig. S5** ANCOM analysis comparing the relative abundance of Planctomycetota in different mangrove sites (KS, PL, SB, MB).

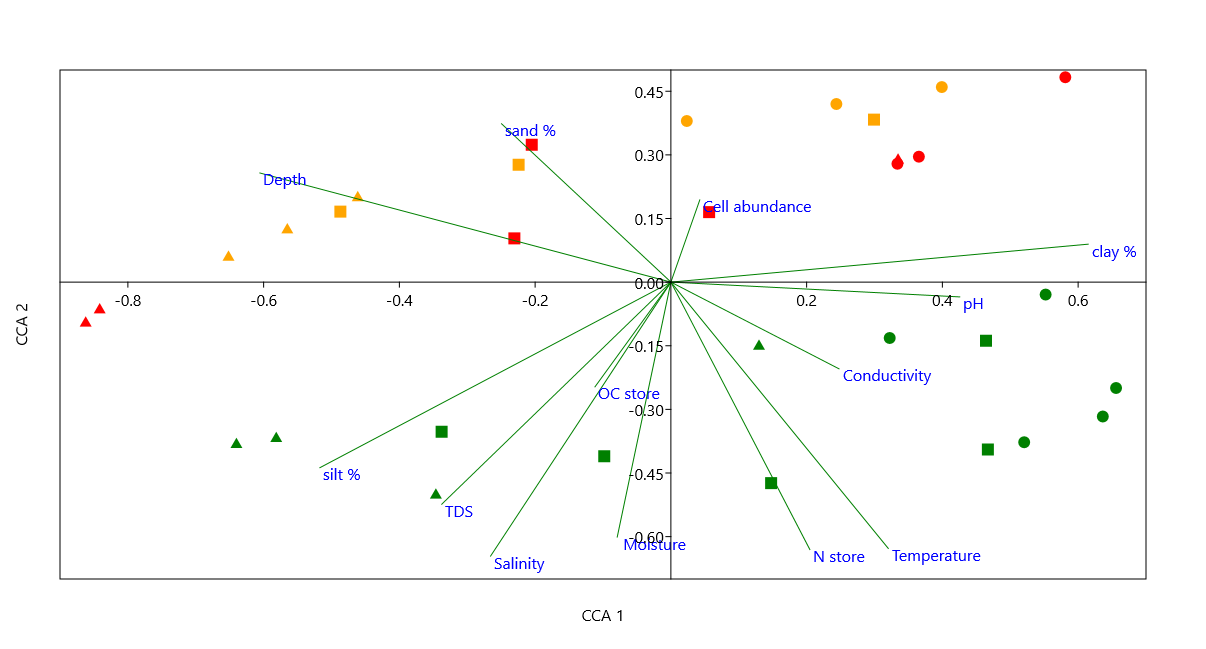

**Fig. S6** Canonical correspondence analysis (CCA) of microbial composition across sites, depths and environmental factors.

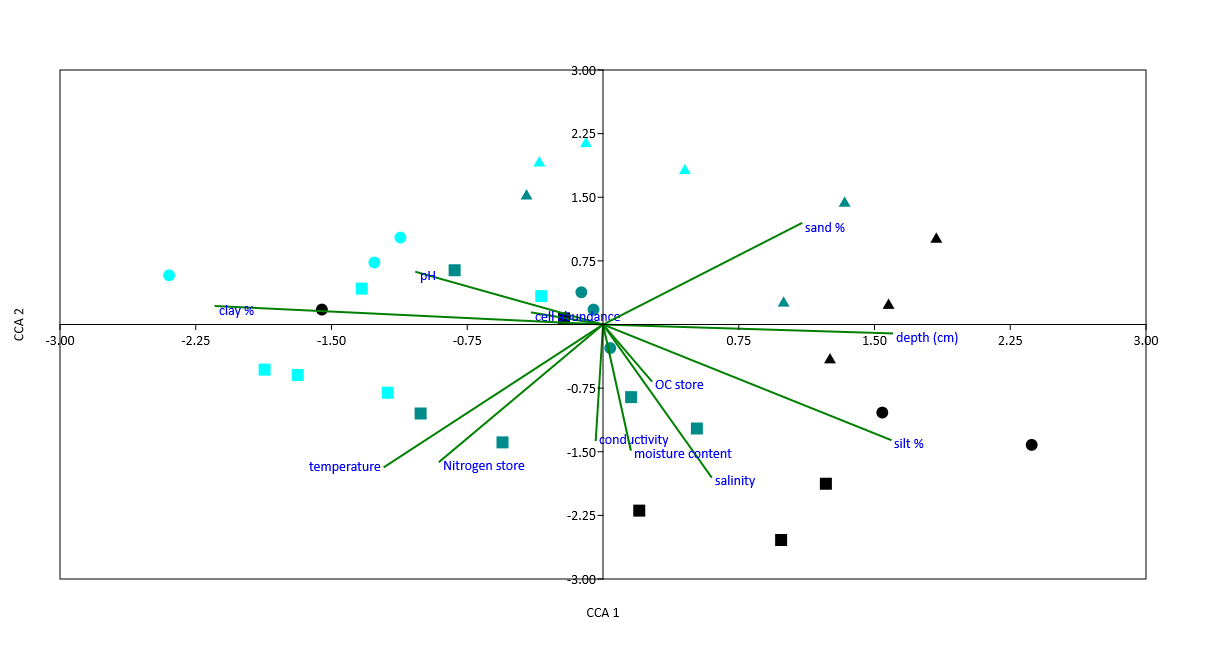

**Fig. S7** CCA of predicted microbial functions (KEGG) across sites, depths and environmental factors.

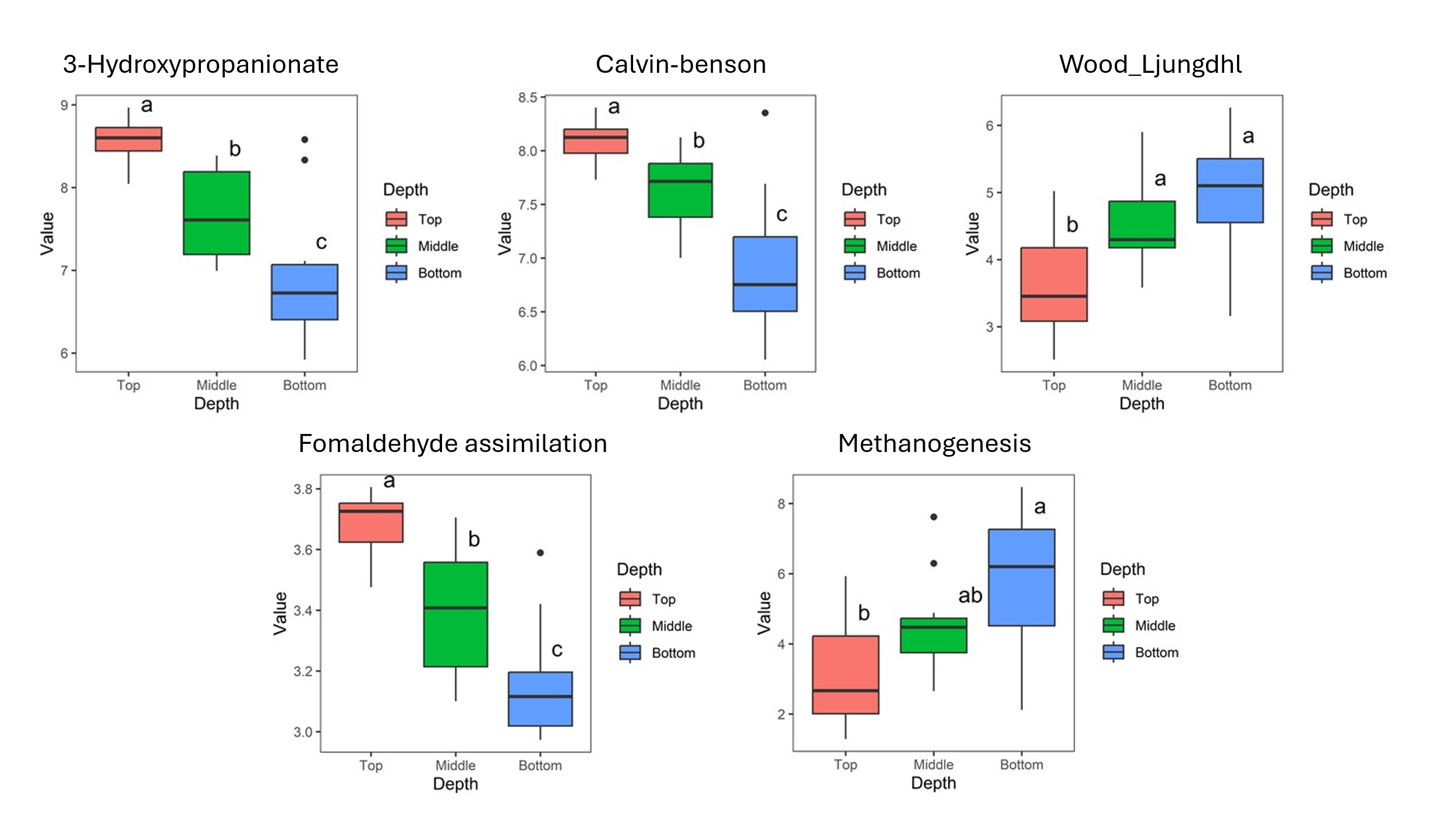

**Fig. S8** Relative abundance of genes related to carbon fixation, methane oxidation, and methanogenesis across sediment depths (top, middle, bottom).

**Table S4** Network analysis of the top twelve microbial phyla based on node characteristics.

| **Phylum** | **Eccentricity** | **Closeness centrality** | **Harmonic closeness centrality** | **Betweenness centrality** |
| --- | --- | --- | --- | --- |
| Latescibacterota | 3 | 0.715789 | 0.806373 | 111.784455 |
| Proteobacteria | 3 | 0.73913 | 0.828431 | 98.556091 |
| Chloroflexi | 3 | 0.693878 | 0.789216 | 88.830653 |
| Desulfobacterota | 3 | 0.571429 | 0.639706 | 84.643077 |
| Archaea_unclassified | 3 | 0.708333 | 0.79902 | 77.081277 |
| Hydrogenedentes | 3 | 0.693878 | 0.784314 | 63.716703 |
| Planctomycetota | 3 | 0.641509 | 0.730392 | 63.714603 |
| Actinobacteriota | 2 | 0.747253 | 0.830882 | 59.629886 |
| Caldatribacteriota | 3 | 0.535433 | 0.585784 | 48.336642 |
| Bacteroidota | 3 | 0.715789 | 0.806373 | 46.803547 |
| Verrucomicrobiota | 3 | 0.647619 | 0.732843 | 44.65758 |
| Dependentiae | 3 | 0.596491 | 0.671569 | 40.981258 |
| Spirochaetota | 3 | 0.747253 | 0.835784 | 40.508097 |
| Asgardarchaeota | 3 | 0.647619 | 0.737745 | 38.211832 |
| Crenarchaeota | 3 | 0.701031 | 0.801471 | 33.816655 |
| Thermoplasmatota | 3 | 0.686869 | 0.781863 | 32.464876 |
| Campilobacterota | 3 | 0.515152 | 0.563725 | 31.061765 |
| Gemmatimonadota | 3 | 0.693878 | 0.789216 | 30.775053 |
| NB1-j | 3 | 0.731183 | 0.82598 | 29.610725 |

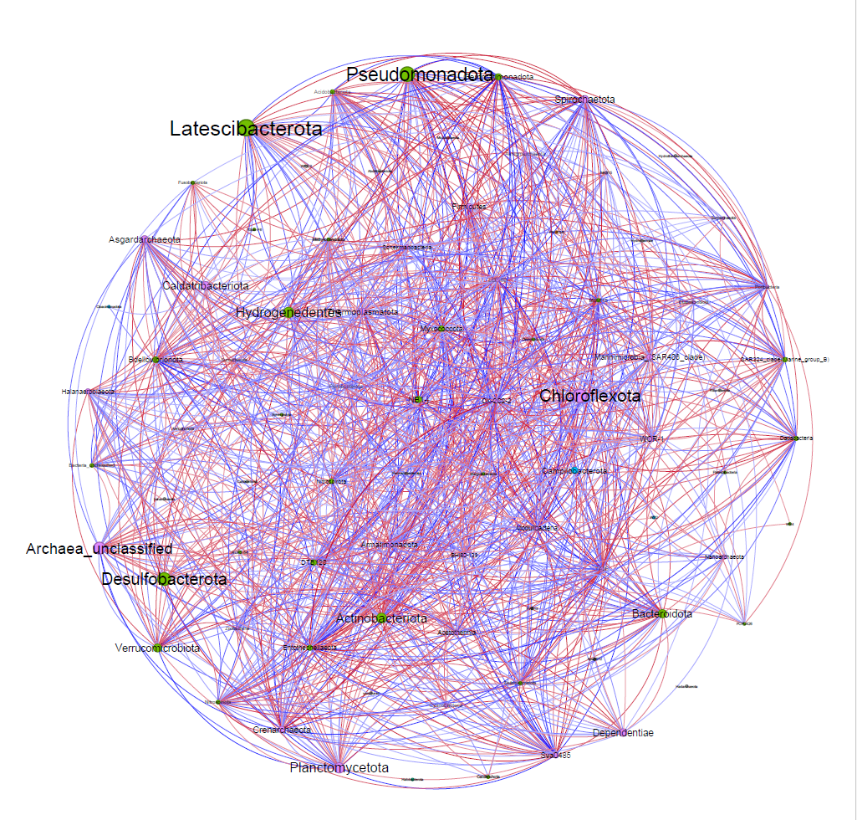

**Fig. S9** Co-occurrence network analysis showing microbial interactions in mangrove sediments.
